# Joint Learning of Node Semantics and Graph Topology using a Transformer in the sparse network regime

**DOI:** 10.1101/2023.12.05.570178

**Authors:** Aidyn Ubingazhibov, David Gomez-Cabrero, Narsis A. Kiani, Jesper Tegner

## Abstract

The human interactome is a valuable tool for unraveling disease mechanisms, advancing precision medicine, facilitating drug discovery, and identifying biomarkers. Yet, current interactomes are incomplete, in part due to limited experimental coverage. Therefore, augmenting the human interactome by predicting missing links in the Protein-Protein interaction network (PPI), is a core challenge for precision medicine. This study proposes an end-to-end trainable transformer-based neural network for enhanced aggregation of Gene Ontology (GO) terms features. We augment the model’s predictive capabilities by incorporating semantic anc2vec features, complementing the structural node2vec embeddings specifically designed for sparse PPIs. As a result, by integrating semantic and graph features, we demonstrate superior performance in link prediction.

## 1 Introduction

Protein-Protein Interactions (PPIs) underpin many biological processes, with over 80% of proteins engaging in complex formations. These interactions weave an intricate web referred to as Protein-Protein Interaction Networks within an organism. Despite the importance of these networks to understanding biological systems, the complete annotation of these interactions remains elusive, primarily due to the constraints imposed by experimental techniques. This gap in knowledge raises the pressing issue of link prediction, a problem that has stimulated significant interest in computational biology and machine learning communities [9]. The challenge of link prediction is accentuated when considering the phenomenon of the “dark proteome”. This term refers to the vast uncharted regions of the protein universe yet to be fully explored or understood. The absence of comprehensive information about these proteins and their potential interactions contributes to the difficulty of comprehensively mapping PPI networks. In response to these challenges, a variety of heuristic methods have been proposed, including Adamic-Adar [1], common neighbors, Katz Index [7], and Clustering Coefficient-based Link Prediction (CCLP) [15]. These approaches have demonstrated broad applicability and robustness [5, 8]. Learning-based methodologies have also been proposed, whereby nodes are mapped into an embedding space via unsupervised methods. The resultant node features are task-independent, with the similarity between node embeddings indicating their similarity within the network. Various random walk-based techniques such as node2vec [6], HOPE [10], and Walklets [12] have proven effective for numerous downstream tasks, including link prediction.

While these methods generate structural features for the nodes in the graph, they do not consider the semantic aspect of the nodes. Since semantic features can offer important indications of link presence, this constitutes a significant gap in the current methodologies. In this regard, anc2vec [3] introduced a novel approach, generating feature embeddings for Gene Ontology (GO) terms using neural networks. This technique captures ontological uniqueness, ancestor hierarchy, and sub-ontology membership, augmenting protein representation beyond mere structural attributes. However, this method’s efficacy is limited when protein gene ontologies are unavailable.

In our work, we further advance this field by improving the anc2vec method and integrating the produced semantic features with structural node2vec. We train a transformer-based neural network on anc2vec features to predict protein interactions, providing an enhanced protein representation that can effectively complement structural node2vec features.

## 2 Problem formulation

The problem of link prediction in protein-protein interaction networks can be formalised as follows: given a graph *G* = (*V, E*) where *V* represents the set of proteins and *E* represents the set of known interactions between proteins, the goal is to predict the set of missing interactions *E′* ⊆ (*V* × *V*) \ *E*.

## 3 Method

GO term features can be effectively represented using anc2vec embeddings, demonstrating state-of-the-art performance among other GO term-based approaches in various down-stream tasks, including link prediction. They obtain protein features by summing the corresponding anc2vec GO term embeddings. In contrast, our approach involves learning a function of the GO terms that yields an enriched protein feature representation. Figure 1 depicts an overview of the model. The number of GO terms associated with proteins can vary significantly. To accommodate the relationships between the anc2vec features of these terms, we propose the utilization of a transformer encoder [14] for learning protein representations. Given the lack of specific ordering of these GO terms, we have excluded positional encoding in our model. To implement this approach, we extract the anc2vec features of each protein’s GO terms and pass them into a 2-layer transformer encoder. We then use global average pooling to obtain protein features. To predict the presence of a link between two proteins, we average their feature representations and use a feed-forward neural network to predict the link score. This method results in an end-to-end trainable network, which we optimize using binary cross-entropy loss.

**Figure 1.**
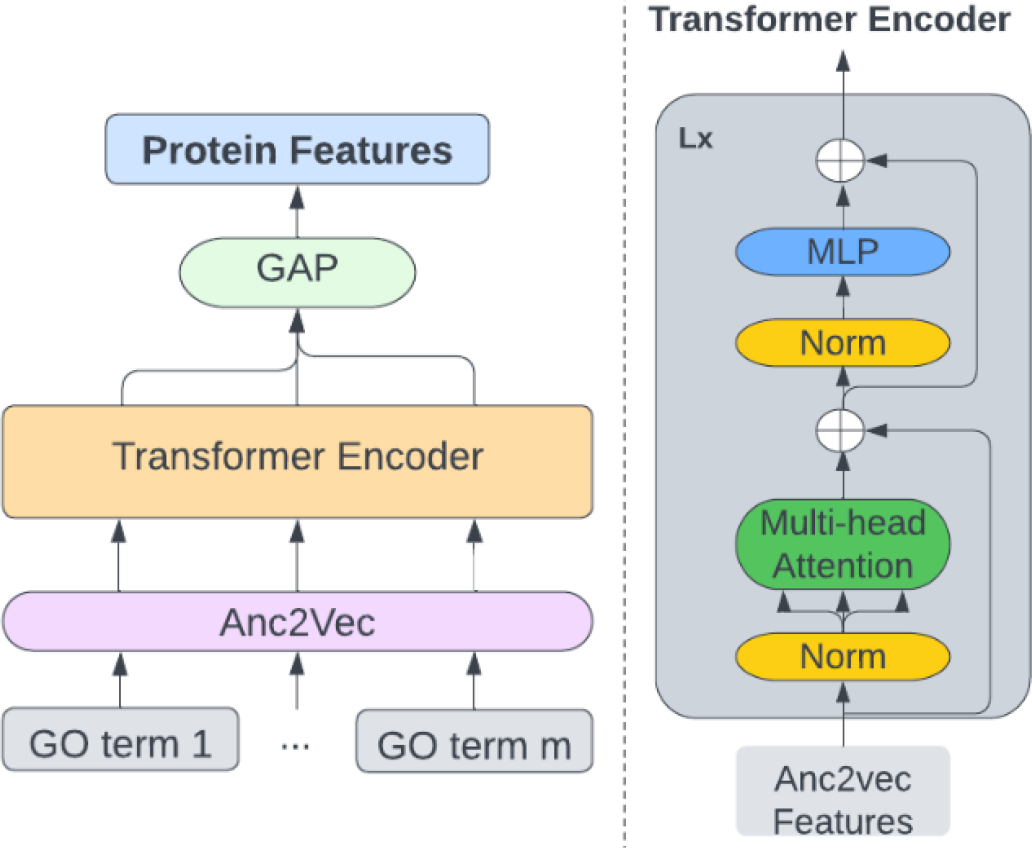
For each protein, anc2vec feature vectors of its GO terms are fed into a standard Transformer Encoder followed by Global Average Pooling (GAP) to produce protein features.

## 4 Semantically enhancing topological features using anc2vec

We focus on *node2vec*, a random walk-based approach for generating structural node features. Return parameter *p* and in-out parameter *q* of the algorithm allow for various exploration strategies of the network, such as Breadth First Search (BFS) or Depth First Search (DFS). It is worth noting that for node classification tasks, it has been shown that *node*2*vec* features can sometimes work as well or even better than actual semantic features for some datasets such as Cora, Pubmed, Reddit, and NELL [2].

In scenarios where the knowledge of existing protein interactions is limited, resulting in a sparse PPI network, the quality of node2vec embeddings tends to deteriorate. Contrarily, the quality of anc2vec features remains unaffected by the extent of known existing links, rendering them a valuable resource for sparse networks. We propose a strategy whereby the structural features procured via node2vec are augmented with semantic features derived from anc2vec, thereby enhancing the prediction of missing links in sparse networks.

More specifically, for a sparse PPI network, we first utilise our transformer model to learn protein features based solely on anc2vec embeddings. We concatenate the learned protein features with node2vec embeddings to generate enriched protein features as shown in Figure 2. This approach ensures that even in the context of limited link information, our method can leverage the strengths of both structural and semantic features, thereby significantly improving the quality of link prediction. The subsequent stages of the process align with the previously described pipeline, highlighting the flexibility and adaptability of this approach.

**Figure 2.**
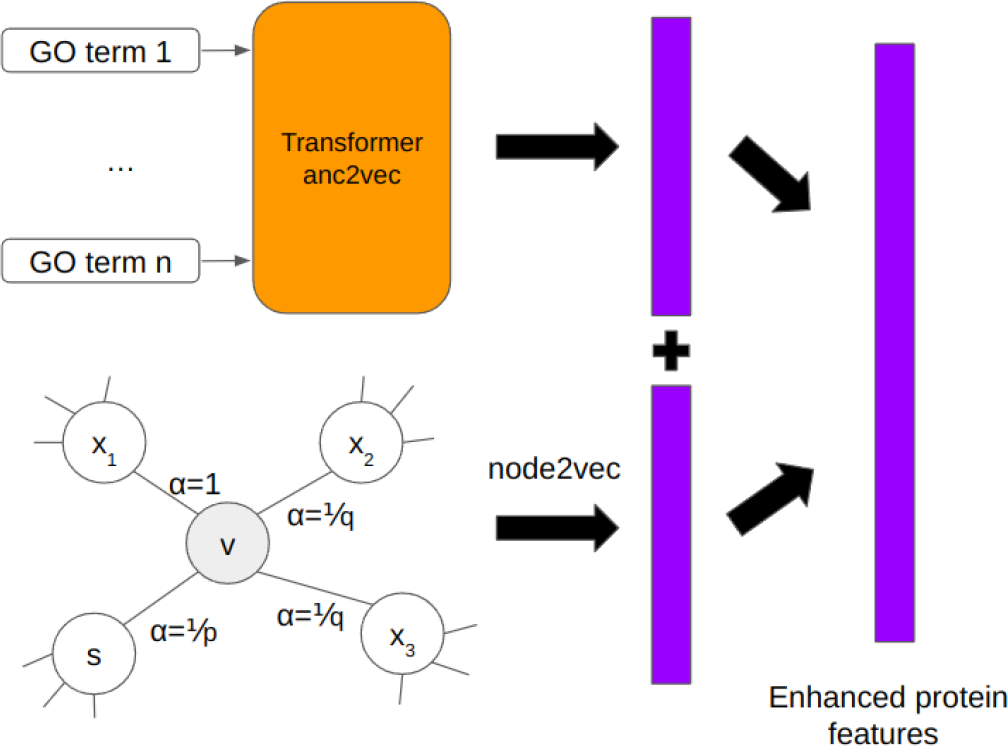
In the context of a sparse network, we generate node2vec features and combine them with anc2vec features produced from a pre-trained transformer model. The figure illustrates the feature extraction process for a specific node ‘v’. In the node2vec representation, ‘v’ is the current node, while ‘s’ is the previous node. ‘p’ and ‘q’ are transition probabilities.

## 5 Datasets

To construct PPI networks, we leveraged STRING database [13], applying an interaction score threshold larger than 0.5. Our experiments were confined to three species: human, mouse, and yeast, which comprise 19354, 21291, and 6574 nodes (proteins), respectively. However, it is worth noting that while mapping the proteins with anc2vec features, specific nodes without GO terms mappings were filtered out.

## 6 Implementation details

For all our experiments, we use Adam optimizer with an initial learning rate of 3 · 10^*−*4^. We train the models for 15 epochs and choose the model based on validation performance. Anc2vec and node2vec both produce 200-dimensional features. The hyperparameters of all of our models (including node2vec) were tuned on the validation set. For our experiments with neural networks and transformers, we use PyTorch [11] [4].

## 7 Experiments

We use the area under the receiver operating characteristic curve (AUROC) as our metric. Each experiment was run six times, and their means and standard deviations were reported to account for variability in the splits and training process.

### 7.1 Data Partitioning

To conduct a comparative analysis between the original anc2vec method and our proposed approach, we adhere to standard practice by partitioning the data as follows: We randomly sample 10% and 20% of the positive edges (without replacement) for validation and testing sets, respectively. The remaining positive edges form the training set. A slightly different strategy is employed when testing extremely sparse networks, with only 10-20% of the existing links available as a training set. In these instances, we randomly sample 10% of the positive (non-training) edges for the validation set with all remaining links serving as the testing edges. We ensure a balanced dataset for every experiment by sampling an equal number of negative and positive edges for each set. This practice mitigates the risk of model bias and helps ensure the generalisability of our results. It’s important to note that, to prevent data leakage, the node2vec embeddings are exclusively generated using the training links.

### 7.2 Comparative Analysis

Table 1 showcases the superiority of our approach in learning protein representation using a transformer encoder over the conventional sum aggregator used in anc2vec [3]. Furthermore, we compare link prediction on protein features derived using our methodology against node2vec embeddings when the PPI’s link knowledge is minimal (10% of data). Table 2 illustrates a sharp decline in performance when deriving node2vec features from highly sparse PPIs. To counter this, we semantically augment the structural features obtained via node2vec with protein features generated by our transformer model for sparse PPIs. The results, displayed in Table 3, clearly indicate a significant improvement in performance using the combined features. It’s worth highlighting that when the entire dataset is available, pure node2vec-based link prediction outperforms the anc2vec-based method, underscoring the greater importance of structural features in indicating link presence.

**Table 1:**
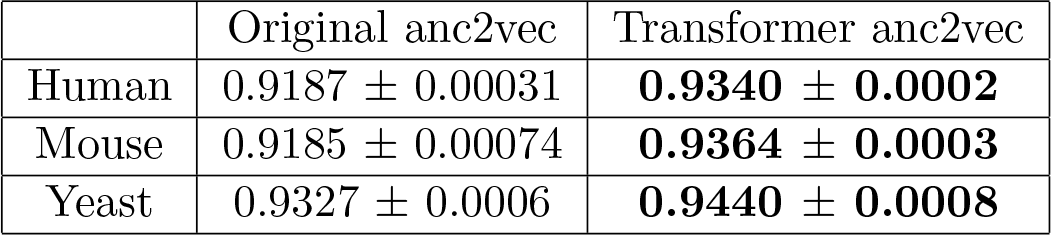
Link prediction using original anc2vec and our approach.

**Table 2:** Link prediction using node2vec features compared against node2vec produced from only 10% data and anc2vec using our approach trained on 10% of the links.

|  | node2vec<br>(full) | node2vec<br>(10%) | Transformer anc2vec<br>(10%) |
| --- | --- | --- | --- |
| Human | $0.9533 \pm 0.0005$ | $0.8742 \pm 0.0008$ | $0.9068 \pm 0.0001$ |
| Mouse | $0.9568 \pm 0.0004$ | $0.8742 \pm 0.0015$ | $0.8979 \pm 0.0008$ |
| Yeast | $0.9617 \pm 0.0007$ | $0.8713 \pm 0.0009$ | $0.8998 \pm 0.0014$ |

**Table 3:** Link prediction on 20% data using node2vec alone, anc2vec alone, and anc2vec combined with node2vec.

|  | Transformer anc2vec | node2vec | combined |
| --- | --- | --- | --- |
| Human | $0.9226 \pm 0.0007$ | $0.9249 \pm 0.0007$ | **$0.9343 \pm 0.0007$** |
| Mouse | $0.9229 \pm 0.0007$ | $0.9198 \pm 0.0006$ | **$0.9344 \pm 0.0007$** |
| Yeast | $0.9131 \pm 0.001$ | $0.9146 \pm 0.0004$ | **$0.9238 \pm 0.0006$** |

## 8 Conclusion

This study contributes to the link prediction field in protein-protein interaction networks, addressing the challenge of sparse data. We proposed a novel method that enhances protein representation by leveraging transformers to learn from Gene Ontology (GO) terms, thereby strengthening the semantic richness of the protein representations. As indicated by the empirical results, our findings demonstrate that our approach outperforms the conventional sum aggregator method, optimizing GO terms-based link prediction. Moreover, our methodology effectively complements the structural features of node2vec, especially when confronted with sparse PPI networks. This dual advantage contributes to improved prediction accuracy and offers resilience against data sparsity.

In conclusion, our approach presents a robust, adaptable, and effective solution for link prediction in PPI networks, addressing the critical limitations of existing methods and providing a solid foundation for future research in this area. Our attempts to employ Graph Autoencoders for link prediction, with node features initialized using the original sum-based anc2vec embeddings, produced unsatisfactory results. One likely factor behind this outcome could be the intricate process of propagating protein features with shared Gene Ontology (GO) terms using Graph Neural Networks (GNNs). This observation underscores the potential difficulties of using GNNs for feature propagation, especially in the context of shared GO terms. We defer the investigation of Graph Neural Networks on anc2Vec features, including our transformer-based anc2vec features, to future research work.

